# Patterns of Treatment in Patients with Heart Failure with Preserved Ejection Fraction: the REasons for Geographic And Racial Differences in Stroke study

**DOI:** 10.1101/510370

**Authors:** Bharat Poudel, Matthew S. Loop, Todd M. Brown, Raegan W. Durant, Monika M. Safford, Parag Goyal, Ligong Chen, Emily B. Levitan

**Author notes:** Address for correspondence: Emily B. Levitan, ScD, University of Alabama at Birmingham, Department of Epidemiology, 1720 2^nd^ Ave S, RPHB 220, Birmingham, AL 35294-0022, USA.

## Abstract

**Purpose:** We described medication use patterns among REasons for Geographic And Racial Differences in Stroke (REGARDS) participants hospitalized for heart failure with preserved ejection fraction (HFpEF) (152 hospitalizations, 101 unique individuals).

**Methods:** Medication data were obtained from medical record review and Medicare Part D pharmacy claims. We compared discharge medication prescriptions between patients with and without chronic kidney disease (CKD), coronary heart disease (CHD), chronic obstructive pulmonary disease (COPD), and diabetes.

**Results:** The mean age was 74.8 years, 53.3% were black and 73.7% were female. Hypertension (97.2%), diabetes (65.1%), COPD (51.3%), CKD (41.1%) and history of CHD (60.9%) were common. On admission and discharge, respectively, beta-blockers (66.4%, 72.7%), angiotensin converting enzyme inhibitors or angiotensin receptor blockers (42.8%, 51.7%), diuretics (61.2%, 80.9%), loop diuretics (55.9%, 78.3%), calcium channel blockers (41.0%, 41.2%) and statins (44.7%, 50.3%) were commonly used. Spironolactone, digoxin, hydralazine plus isosorbide dinitrate (HISDN), isosorbide dinitrate alone and aldosterone receptor antagonists were used by <20%. For each medication, prescriptions were more common at discharge than admission. Many participants did not have Medicare claims for filled prescriptions in the year following discharge. A higher percentage of patients with versus without CKD, CHD, and diabetes had discharge prescriptions statins. Participants with CKD were also more likely to receive prescriptions for HISDN.

**Conclusion:** Beta-blockers and diuretics were commonly prescribed at admission and discharge among HFpEF, but pharmacy claims for these medications within one-year were substantially less common. The comorbidities CHD, CKD, and diabetes were associated with prescriptions of statins at discharge.

## INTRODUCTION

Heart failure with preserved ejection fraction (HFpEF) is a clinical condition with signs and symptoms of heart failure (HF) with normal or near normal left ventricular ejection fraction (≥50%) [1]. The proportion of HF hospitalizations characterized by preserved ejection fraction increased from 33% to 39% during 2005 to 2010 [2, 3]. Clinical trials of therapies targeting the renin-angiotensin-aldosterone system and the adrenergic nervous system have not shown benefit in patients with HFpEF [4]. The Candesartan in Heart Failure-Assessment of Reduction in Mortality and Morbidity (CHARM) and Irbesartan in Patients with Heart Failure and Preserved Ejection Fraction (I-PRESERVE) studies show that neither candesartan nor irbesartan reduced cardiovascular mortality or hospital admissions of patients with HFpEF [5]. Similarly, the result from Treatment of Preserved Cardiac Function Heart Failure with an Aldosterone Antagonist (TOPCAT) trial showed no difference in total deaths and hospitalizations in HFpEF patients randomized to take spironolactone versus placebo [6]. Currently, there are no medications specifically indicated for HFpEF apart from diuretics to treat signs and symptoms of congestion.

The concomitant presence of cardiovascular and non-cardiovascular comorbidities makes diagnosing HFpEF difficult and can complicate management decisions related to drug therapy. Despite no known evidence of benefit of treatment on HFpEF-related outcomes, beta-blockers, angiotensin converting enzymes or angiotensin receptor blockers (ACEIs/ARBs), calcium channel blockers (CCBs), statins and other medications have been commonly used in HFpEF patients to treat associated comorbid conditions [7] as suggested by guidelines for the management of heart failure [8] [9] [10]. The treatment guidelines for HFpEF focus on controlling physiologic factors that adversely affect ventricular diastolic function, which includes blood pressure, heart rate, dyspnea, volume overload, blood volume and myocardial ischemia.

The medication treatment patterns among HFpEF patients have largely been described in single-site studies, homogeneous community based cohorts, or in clinical trials with restrictive eligibility criteria. We aimed to describe medication patterns among a sample of participants that spanned across the entire contiguous United States. This study provides a description of medications being taken upon hospital admission, discharge prescriptions, and prescription fills within one-year post-hospitalization among the REasons for Geographic and Racial Differences in Stroke (REGARDS) study participants that were hospitalized for HFpEF. We also examined the associations between presence of chronic kidney disease (CKD), coronary heart disease (CHD), COPD, and diabetes with medication prescriptions at discharge for HFpEF. We focused on these comorbid conditions because they could have influenced the prescription of the medications of interest.

## METHODS

### Study population

REGARDS is a national cohort study that enrolled 30,239 non-Hispanic white and black adults ≥45 years of age between January 2003 and October 2007 across the 48 contiguous US states. Adults in the Southeastern states and blacks were oversampled in order to study the variation in stroke mortality across geographical regions and race [11]. The data collected from the REGARDS participants were linked to Medicare claims using participants’ social security numbers, date of birth and sex as previously described [12]. Approximately two thirds of participants (n =20,403) had Medicare linkage at some point between 2000 and 2012; approximately one third (n =10,839) had Medicare fee-for-service coverage on the date of study enrollment. Among REGARDS participants ≥65 years of age, 80% had data linked to Medicare and 64% had fee-for-service coverage at study enrollment. In this cross-sectional study, we examined patterns of pharmacologic treatment of patients hospitalized with HFpEF included in the REGARDS cohort and REGARDS-Medicare linkage from 2006 to 2011. Hospitalizations prior to 2006 were not considered because the Medicare Part D prescription drug benefit began in 2006. We identified HF hospitalizations that occurred during periods when successfully linked participants had Medicare hospital, medical and prescription drug claims available. HF hospitalizations with ejection fraction ≥50% or a qualitative report of normal ejection fraction documented in the medical records or a Medicare ICD-9 diagnosis specifying isolated diastolic dysfunction (428.3×) during the current hospitalization or, in the absence of information about ejection fraction during the current hospitalization, a history of normal ejection fraction during a prior medical encounter documented in the medical records were considered HFpEF (n =152 hospitalizations). Participants could contribute more than one hospitalization. Of 600 identified hospitalizations (314 unique patients) for HF linked to Medicare, medical records for 400 hospitalizations were retrieved and 344 had discharged medication lists available. Of the 344 hospitalizations, 147 had HF with reduced ejection fraction (HFrEF), 45 had an unknown ejection fraction, and 152 (101 unique patients) had HFpEF.

### Medications

The admission and discharge medication lists were obtained from a review of medical records from HFpEF hospitalizations. Prescription fills in the 365 days following discharge were obtained from Medicare Part D claims for the participants. Based on the available guidelines on treatment of HFrEF and HFpEF, we focused on beta-blockers, beta-blockers specifically indicated for HFrEF (carvedilol, sustained-release metoprolol succinate, bisoprolol), angiotensin converting enzyme inhibitors (ACEIs) or angiotensin receptor blockers (ARBs), all diuretics and loop diuretics which are recommended for HFpEF, hydralazine in combination with isosorbide dinitrate (HISDN), isosorbide dinitrate without hydralazine, aldosterone receptor antagonists, calcium channel blockers (CCBs), statins, digoxin and spironolactone.

### Covariates

Information on age, race, gender, education, and income was collected during the REGARDS baseline interview. Systolic and diastolic blood pressure (BP) and body mass index (BMI) were measured at the time of hospitalization and from baseline in-home visit, respectively. We determined the presence of comorbidities from documentation on medical records at the time of HFpEF hospitalization. Comorbidities included asthma, COPD, hypotension, atrial fibrillation (AF), peripheral vascular disease and stroke, history of hypertension, diabetes, CKD, dialysis, implanted cardiac devices, myocardial infarction (MI), history of coronary heart disease (CHD), and CHD events within past year, which were also collected at the time of hospitalization. Hospitalization within past year and outpatient clinic visits in the week after discharge were detected using Medicare claims.

### Statistical Analysis

The population characteristics were described using percentage (%) for categorical variables and mean ± standard deviation (SD) for continuous variables. We calculated the percentage of hospitalizations with the medications on admission, at discharge and with claims for prescription fills in the 365 days post-discharge. REGARDS participants with HFpEF were stratified based on the comorbid conditions CKD, CHD, COPD, and diabetes to compare medication prescriptions at discharge. Prevalence of discharge medication prescriptions for those with and without the comorbidities compared using generalized estimating equation Poisson models with robust variance estimates to calculate prevalence ratios, accounting for individual participants contributing multiple hospitalizations. We did not examine the association of comorbidities with use of isosorbide dinitrate without hydralazine because of the small number of patients who used this agent. To account for missing data, we used the fully conditional specification (FCS) method of multiple imputation to create 20 imputed datasets. Analyses were conducted in each dataset; point estimates and standard errors were combined across datasets [13]. All statistical analyses were performed using SAS version 9.4. Software (SAS Institute, Inc., Cary, North Carolina).

### Ethical Considerations

REGARDS study participants provided informed consent for participation in research, including linkage with Medicare claims, and signed medical record release forms allowing study investigators to retrieve hospitalization records for research. The Centers for Medicare and Medicaid Services Privacy Board and the University of Alabama at Birmingham Institutional Review Board approved this research.

## RESULTS

Among 600 hospitalizations, we were able to retrieve medical record and link Medicare prescription drug claims for 152 hospitalizations (101 unique patients) for HF with ejection fraction ≥ 50%, qualitative “normal” ejection fraction, or diagnosis code specifying diastolic heart dysfunction without systolic dysfunction. The mean age was 74.8 ± 8.5 years (**Table 1**). More than half of the participants with HFpEF were black, and 73.7% were female. The mean ± SD of systolic and diastolic blood pressure of the participants at the time of hospitalization were 151 ± 35 mmHg and 75 ± 22 mmHg respectively. More than 55% of the participants were obese and nearly 35% of the participants had less than a high school of education. Approximately 62% of the participants had annual income less than $20,000. Most of the patients hospitalized for HFpEF had a history of hypertension (97.2%), diabetes (65.1%), history of coronary heart disease (60.9%), and COPD (51.3%) while 38.9% and 41.1% had AF and CKD, respectively.

**Table 1:** Participant characteristics and comorbidities present in REGARDS participants with HFpEF (N=152 hospitalizations).

| **Demographic characteristics** | **% or Mean ± SD** |
| --- | --- |
| Age | 74.8 ± 8.5 |
| $\leq 65$ years | 13.2 |
| 65 – 69 years | 15.1 |
| 70 – 74 years | 21.7 |
| 75 – 79 years | 21.7 |
| 80 – 84 years | 13.2 |
| $\geq 85$ years | 15.1 |
| Race |  |
| Whites | 46.7 |
| African Americans | 53.3 |
| Gender |  |
| Male | 26.3 |
| Female | 73.7 |
| Blood Pressure (BP) at the time of hospitalization |  |
| Systolic BP, mm Hg | 151 ± 35 |
| Diastolic BP, mm Hg | 75 ± 22 |
| Body Mass Index (kg/m<sup>2</sup>) | 32.8 ± 7.5 |
| Underweight (<18.5) | 7.9 |
| Normal (18.5 – 24.9) | 11.8 |
| Overweight (25.0 – 29.9) | 23.0 |
| Obese ($\geq 30.0$) | 55.9 |
| Education |  |
| Less than High School | 35.5 |
| High School Graduate | 32.9 |
| Some College | 21.7 |
| College Graduate and above | 9.9 |
| Annual Income |  |
| < \$ 20,000 | 61.8 |
| \$20,000 – \$ 34,999 | 20.4 |
| \$ 35,000 - \$ 74,999 | 5.9 |
| ≥ \$ 75,000 | 2.0 |
| Declined to report | 9.9 |
| **Comorbidities** |  |
| Asthma | 9.9 |
| COPD | 51.3 |
| AF | 38.9 |
| Peripheral Vascular Disease | 12.2 |
| Stroke | 23.7 |
| History of HTN | 97.2 |
| Diabetes | 65.1 |
| CKD | 41.1 |
| Dialysis | 17.2 |
| Implanted cardiac device | 19.7 |
| MI | 30.6 |
| History of CHD | 60.9 |
| CHD event within past year | 7.3 |
| Hospitalization within past year | 71.1 |
| **Characteristics of Hospitalization** |  |
| Ejection fraction | 55 (55 – 60)* |
| Hypotension | 13.0 |
| COPD contributed to hospital admission | 13.2 |
| Outpatient clinical visit in week after discharge | 23.0 |

COPD= Chronic Obstructive Pulmonary Disease, AF= Atrial Fibrillation, HTN= Hypertension, CKD= Chronic kidney disease, MI= Myocardial Infraction, CHD= Coronary heart disease
\*Expressed as median (25<sup>th</sup> - 75<sup>th</sup> percentile).

Among hospitalizations for HFpEF, at admission 66.4% had prescriptions for beta-blockers, 42.8% had prescriptions for ACEIs/ARBs, 61.2% had prescriptions for diuretics, 44.7% had prescriptions for statins, and 41% had prescriptions for CCBs (Table 2). Only 20.4% of patients had prescriptions for beta-blockers with guideline indications for systolic dysfunction (HFrEF), 15.1% had HISDN and 6.6% had aldosterone receptor antagonists and spironolactone at admission. At discharge, 72.7% had prescriptions for beta-blockers, and nearly 80.0% had prescriptions for diuretics including loop diuretics. The majority of patients filled prescriptions for diuretics (73.0%), loop diuretics (70.4%), and beta-blockers (58.6%) within one year of hospital discharge. A smaller percentage of patients had prescription claims within one year for CCBs (10.5%) compared to their prescriptions at admission (41.0%) and discharge (41.2%). The use of spironolactone (6.6%), digoxin (17.6%), HISDN (15.1%), isosorbide dinitrate without hydralazine (1.3%) and aldosterone receptor antagonists (6.6%) at admission was relatively lower. The prevalence of discharge prescriptions and pharmacy claims within one year after discharge for these medications were lowest of the therapies studied.

**Table 2:**
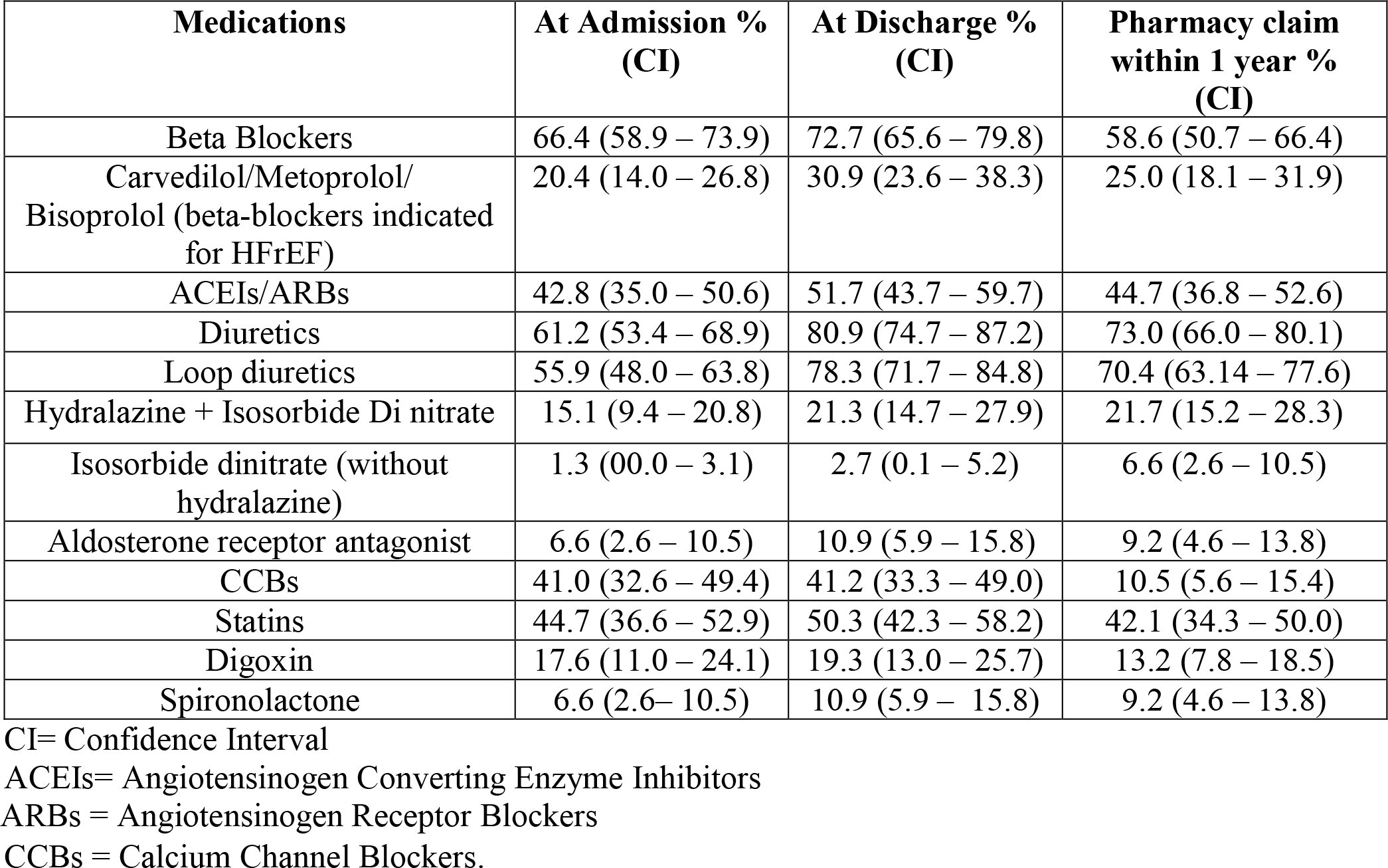
Medication patterns among REGARDS participants with HFpEF (N=152 hospitalizations)

Few differences were present in the prescriptions among patients with and without presence of comorbidities (Table 3). Discharge prescriptions for HISDN and statins were more common among patients with versus without CKD (prevalence ratio 2.3, CI [1.1-4.9] and 1.6 [1.1-2.3] respectively). Prescriptions for statins were more common among patients with CHD (1.9 [1.2-2.8]) and diabetes (2.0 [1.2-3.3]) compared to their counterparts without these comorbidities.

**Table 3:** Comorbidities and medications prescribed at discharge among REGARDS participants with HFpEF

| Medications | Comorbidities |  |  |  |  |  |  |  |  |  |  |  |
| --- | --- | --- | --- | --- | --- | --- | --- | --- | --- | --- | --- | --- |
|  | CKD |  |  | CHD |  |  | COPD |  |  | Diabetes |  |  |
|  | Yes | No | PR (CI) | Yes | No | PR (CI) | Yes | No | PR (CI) | Yes | No | PR (CI) |
| Beta-Blockers | 76.0 | 70.4 | 1.1 (0.9-1.4) | 72.4 | 73.1 | 1.0 (0.8-1.2) | 64.1 | 81.8 | 0.8 (0.6-1.0) | 70.2 | 77.4 | 0.9 (0.7-1.1) |
| Carvedilol/Metoprolol/<br>Bisoprolol (beta-blockers<br>indicated for HFrEF) | 28.0 | 33.0 | 0.8 (0.4-1.6) | 30.1 | 32.3 | 0.9 (0.5-1.6) | 30.8 | 31.1 | 1.0 (0.6-1.7) | 33.3 | 26.4 | 1.3 (0.7-2.2) |
| ACEIs/ARBs | 44.2 | 56.9 | 0.8 (0.5-1.1) | 52.8 | 49.9 | 1.1 (0.7-1.5) | 47.4 | 56.1 | 0.8 (0.6-1.2) | 50.1 | 54.7 | 0.9 (0.7-1.2) |
| Diuretics | 71.2 | 87.7 | 0.8 (0.7-1.0) | 83.8 | 76.5 | 1.1 (0.9-1.3) | 82.1 | 79.7 | 1.0 (0.9-1.2) | 82.8 | 77.4 | 1.1 (0.9-1.3) |
| Loop Diuretics | 68.8 | 85.2 | 0.8 (0.6-1.0) | 81.5 | 73.3 | 1.1 (0.9-1.3) | 82.1 | 74.3 | 1.1 (0.9-1.4) | 80.8 | 73.6 | 1.1 (0.9-1.4) |
| Hydralazine with<br>Isosorbide dinitrate | 31.8 | 14.0 | 2.3 (1.1-4.9) | 22.3 | 19.8 | 1.1 (0.6-2.2) | 25.6 | 16.8 | 1.5 (0.8-3.1) | 23.6 | 17.0 | 1.4 (0.7-2.8) |
| Aldosterone receptor<br>antagonist | 5.2 | 14.8 | 0.3 (0.1-1.1) | 11.6 | 9.7 | 1.2 (0.4-3.4) | 9.0 | 12.8 | 0.7 (0.2-2.0) | 12.6 | 7.5 | 1.7 (0.5-5.1) |
| CCBs | 45.6 | 38.1 | 1.2 (0.8-1.9) | 43.5 | 37.6 | 1.1 (0.8-1.7) | 41.0 | 41.4 | 1.0 (0.7-1.5) | 37.0 | 49.1 | 0.7 (0.5-1.1) |
| Statins | 65.0 | 39.9 | 1.6 (1.1-2.3) | 61.6 | 32.6 | 1.9 (1.2-2.8) | 50.0 | 50.5 | 1.0 (0.7-1.4) | 61.0 | 30.2 | 2.0 (1.2-3.3) |
| Digoxin | 12.2 | 24.4 | 0.5 (0.2-1.2) | 17.7 | 21.9 | 0.8 (0.3-2.0) | 21.8 | 16.8 | 1.3 (0.6-2.8) | 19.6 | 18.9 | 1.0 (0.4-2.5) |
| Spironolactone | 5.2 | 14.8 | 0.3 (0.1-1.1) | 11.6 | 9.7 | 1.2 (0.4-3.4) | 9.0 | 12.8 | 0.7 (0.2-2.0) | 12.6 | 7.5 | 1.7 (0.5-5.1) |

PR: Prevalence Ratio, CI: Confidence Interval.

## DISCUSSION

This study of REGARDS participants hospitalized for HFpEF demonstrated that patients were likely to be treated with diuretics and medications indicated for HFrEF including beta-blockers and ACEIs/ARBs. The majority were prescribed diuretics and beta-blockers both prior admission and at discharge. For all medications examined, the percentage of participants who were prescribed medications at discharge was greater than at admission, but many participants did not have Medicare claims for filled prescriptions in the year following discharge.

The American College of Cardiology/ American Heart Association (ACC/AHA), European Society of Cardiology and Heart Failure Society of America (HFSA) treatment guidelines recommend controlling factors affecting ventricular diastolic function, which includes controlling blood pressure, heart rate, blood volume, and myocardial ischemia in HFpEF patients [8-10]. These guidelines also recommend the use of diuretics to relieve symptoms due to volume overload [8, 9] and hypertensive patients with concomitant presence of CKD are recommended diuretics and ACEIs/ARBs regardless of diabetes status. Similarly, a thiazide diuretic, ACEIs/ARBs or CCBs are appropriate for hypertensive patients who do not have concomitant kidney disease [14]. Hypertension was most common comorbid condition occurring in 97.2% in the participants of our study. The use beta-blockers, ACEIs/ARBs, diuretics and CCBs at admission, discharge and one-year refill was likely to be in large part intended to treat hypertension. CHD, MI, and AF were also common as were other non-cardiac comorbidities like COPD, diabetes, CKD, and obesity. Stratification of hospitalized HFpEF participants based on presence or absence of CKD, CHD, COPD and diabetes showed that treatment patterns were largely similar irrespective of the comorbidities present except for use of HISDN and statins in participants with CKD and statins in participants with CHD and diabetes where the discharge prescriptions of these medications were higher compared to those who did not have comorbidities.

Several pharmacological therapies including diuretics, beta-blockers, and ACEIs/ARBs have improved outcomes of morbidity and mortality in HFrEF patients in the last two decades, but long term clinical outcomes in patients with HFrEF and HFpEF are still poor [15-17]. Various clinical trials have been conducted to improve the outcomes in HFpEF patients targeting clinical symptoms, exercise capacity, diastolic function and quality of life, but none of them was successful in reducing mortality [17] [18]. The Hong Kong Diastolic Heart Failure Study, which evaluated the use of diuretics (furosemide or thiazide) alone or combined with ACEI (ramipril) or ARB (irbesartan) in 150 patients with LVEF > 45%, showed improvement of symptoms and quality of life when diuretics were used alone, with only slight additional benefits observed when diuretics were combined with ramipril or irbesartan [19]. CHARM-Preserved, I-PRESERVE and PEP-CHF studies did not find evidence that ACEIs (perindopril) and ARBs (candesartan and irbesartan) reduce the risk of hospitalization or death in patients with HFpEF [20];[21];[22]. Owing to the inconclusive results, ACEIs/ARBs and beta-blockers are not considered as guideline-based therapy in patients with HFpEF unless hypertension and other comorbidities like left ventricular hypertrophy (LVH) and atherosclerotic vascular diseases are present [23].

Treatment patterns in patients with HFpEF have been investigated in several study populations [24] [25] [26]. In a study from the Cardiovascular Research Network that included individuals insured by comprehensive health plans, newly diagnosed patients with HFpEF were more likely to be treated with beta-blockers (62.4%), thiazide and loop diuretics (61.4%), statins (48.0%), ACEIs/ARBs (49.6%), and CCBs (33.8%) than patients with HFrEF prior to HF diagnosis [24]. However, use of these treatments were more common among HFrEF patients than HFpEF patients after diagnosis. In OPTIMIZE-HF registry, the use of beta-blockers was evaluated in 7154 HF hospitalized patients (divided into 2 cohorts-left ventricular systolic dysfunction and patients with HFpEF), which showed HFpEF patients were likely to be prescribed diuretics (79.6%), ACEIs/ARBs (64.5%), and statins (31.5%) but had fewer prescriptions of aldosterone antagonist (9.3%) and digoxin (20.4%) at discharge [26]. We observed similar treatment patterns among REGARDS participants with HFpEF. A report from Japanese Cardiac Registry of Heart Failure in Cardiology (JCARE-CARD) showed a higher use CCBs in HFpEF (42.9%) compared to HFrEF (17.1%) patients when discharged [27]. A study conducted in a local public hospital in Hong Kong with 73 HFpEF patients found that loop diuretics (80.8%), ACEIs (65.8%), beta-blockers (58.9%) and CCBs (57.5%) were commonly prescribed for HFpEF management in an outpatient setting [25]. In the Japanese Cardiac Registry of Heart Failure in Cardiology compared to HFrEF, in HFpEF patients, beta-blockers were less likely to be prescribed at discharge [27], and prescription of diuretics at discharge was similar between groups (84% vs 81%). Most hypertensive patients with HFpEF receive beta-blockers, diuretics and ACEIs/ARBs to control high BP [9]; [28].

The results of this study must be considered in light of its limitations. This study includes a small sample of participants with HFpEF. The small sample size also prevented us from conducting models to adjust for any confounders. However, in contrast to the available studies from registries and specific medical centers, the REGARDS study provides more information on demographics and includes a broad range of participants treated in unselected hospitals from across the United States.

In conclusion, beta-blockers, ACEIs/ARBs, statins, CCBs and diuretics (including loop diuretics) were commonly prescribed for HFpEF patients at admission and discharge. The Medicare claims for medication fills after one year was significantly lower as compared to medications prescribed during discharge in HFpEF patients suggesting that many individuals do not take prescribed medications. The discharged prescriptions for HISDN and statins were higher among those with CKD compared to those without CKD. Similarly, the prescriptions for statins were significantly higher in patients with CHD and diabetes compared to patients without those comorbidities.

## Acknowledgements

The authors thank the other investigators, the staff, and the participants of the REGARDS study for their valuable contributions. A full list of participating REGARDS investigators and institutions can be found at http://www.regardsstudy.org.

## Declarations

Availability of data and materials: The data that support this study are available from the REGARDS study (www.regardsstudy.org), but restrictions apply to the availability of these data. The data are not publicly available because they contain information that could compromise research participant privacy and consent.

Ethics approval and consent to participate: This research was conducted in accordance with the Declaration of Helsinki. The REGARDS study was approved by the Institutional Review Boards of the University of Alabama at Birmingham, the University of Vermont, Wake Forest University, and the University of Cincinnati. Participants provided written informed consent and completed medical record release forms authorizing study investigators to retrieve medical records for research purposes. The Institutional Review Board of the University of Alabama at Birmingham approved the current research.

